# Maximum Parsimony Reconciliation in the DTLOR Model

**DOI:** 10.1101/2021.05.23.445352

**Authors:** Jingyi Liu, Ross Mawhorter, Nuo Liu, Santi Santichaivekin, Eliot Bush, Ran Libeskind-Hadas

## Abstract

**Background:** Analyses of microbial evolution often use reconciliation methods. However, the standard duplication-transfer-loss (DTL) model does not account for the fact that species trees are often not fully sampled and thus, from the perspective of reconciliation, a gene family may enter the species tree from the outside. Moreover, within the species tree, genes are often rearranged, causing them to move to new syntenic “regions.”

**Results:** We extend the DTL model to account for two events that commonly arise in the evolution of microbes: evolution occurring outside the sampled species tree and changes in the syntenic regions of genes in the genome due to rearrangement. We describe an efficient algorithm for maximum parsimony reconciliation in this new DTLOR model and then show how it can be extended to account for non-binary gene trees. Finally, we describe preliminary experimental results from the integration of our algorithm into the existing xenoGI tool for reconstructing the histories of genomic islands in closely related bacteria.

**Conclusions:** Reconciliation in the DTLOR model can offer new insights into the evolution of microbes that is not currently possible under the DTL model.

## 1 Background

Microbes occupy a vast range of ecological niches [1]. Understanding how particular species have come to occupy their niches requires us to reconstruct how their genomes have evolved over time.

In a clade of closely related microbes with a known gene and species tree, inferring the genetic history can be done through a process called *reconciliation*. This process maps the gene tree to the species tree, and in doing so implies genetic events that explain the discordance between the two trees. The DTL model considers duplication, horizontal gene transfer, and loss events whereas some models consider a subset of these events (e.g., only duplication and loss) or different types of events (e.g,. independent lineage sorting).

While the DTL model is applicable to evolution in microbes, that model only allows horizontal transfer between species that are part of the species tree. In the analysis of the evolution of microbes in particular, it is quite common that the species tree is not fully sampled. Thus, from the perspective of performing a reconciliation analysis, a gene family may effectively enter the given species tree via transfer from the outside [2, 3].

In this paper we describe the DTLOR model that addresses this issue by extending the DTL model to allow some or all of the evolution of a gene family to occur outside of the given species tree and for transfers events to occur from the outside. To facilitate the recognition of such entry events, the model also keeps track of the *syntenic region* of genes as they evolve in the species tree. Two genes are said to be in the same syntenic region if they share a substantial fraction of core genes in a relatively large window around them and, second, they share a certain amount of similarity among all genes in a smaller window around them [2]. Thus, in addition to duplication, transfer, and loss events, the DTLOR model adds *origin* events to indicate that a gene is transferred from outside of the species tree and *rearrangement* events that account for changes in the syntenic regions of genes on the genome.

In the DTL model, reconciliation is generally performed using a maximum parsimony formulation. A positive cost is associated with each type of event and the objective is to find a reconciliation that minimizes the total cost of the incurred events. Efficient algorithms have been developed for maximum parsimony reconciliation in the DTL model [4–6] and several software tools implement these algorithms [7–10].

In earlier related work, Szöllösi *et al.* [3] proposed an event called “transfer from the dead” to account for gene evolution that occurs outside of the species tree and Jacox *et al.* [10] described an extension of an existing DTL maximum parsimony reconciliation algorithm to compute most parsimonious reconciliations with this additional event. Our work differs in two significant ways from this prior work. First, while “transfer from the dead” allows gene lineages to transfer out and back in of the sampled species trees multiple times, the DTLOR model only permits transfer from the outside under the assumption that the species tree comprises closely-related species and, thus, transfers out of the species tree and back in are considered to be relatively rare. Second, the DTLOR model captures rearrangement events, which are not considered in conjunction with DTL events in previous models. Reconstructing rearrangement events is particularly important in identifying genomic islands in bacteria [2].

In summary, in this paper we extend the DTL model to allow origin (O) and rearrangement (R) events. We give an exact polynomial-time algorithm for maximum parsimony reconciliation in the DTLOR model. Since gene trees are often non-binary due to lack of signal in their sequence data, we show how maximum parsimony reconciliations can be found in fixed-parameter polynomial time for non-binary gene trees. Finally, we describe preliminary results from the integration of the DTLOR MPR algorithm into the xenoGI tool for microbial evolution [2].

## 2 Background and Definitions

### 2.1 Overview of the DTLOR MPR Problem

An instance of the DTLOR reconciliation problem comprises undated species and gene trees, *S* and *G*, respectively; a positive integer syntenic region number associated with each leaf vertex (extant gene) in *G*; and a mapping of the leaves of *G* to the leaves of *S*. The DTLOR model comprises the standard DTL events (speciation, duplication, transfer, and loss; described in detail below) and two additional events called *origin* and *rearrangement*. Each of those five event types has an associated positive cost.

A syntenic region number is a positive integer from the set of syntenic region numbers of the leaves of the gene tree (called an *actual syntenic region*) or the special *unknown syntenic region* symbol *. When a gene vertex is labeled with *, that vertex is assumed to be evolving outside the gene tree. When a gene vertex is assigned an actual syntenic region but its parent has unknown syntenic region, this means that the gene entered the species tree through transfer from outside. This induces an origin event. Rearrangement indicates a change in syntenic region that happens during the course of evolution within the species tree.

The rules for syntenic region numbering are as follows:

1. If a vertex *u* is labeled * and *v* is a child of *u*, then *v* may be labeled with either * or an actual syntenic region number.
2. If a vertex *u* is labeled with an actual syntenic region number then its children must be labeled with actual syntenic region numbers. Note that this implies that any vertex labeled with an actual syntenic region number has the property that all of its descendants are also labeled with actual syntenic region numbers.

The DTL events in this model are analogous to those in the DTL model. O and R events are induced as follows:

1. If a vertex *u* is labeled * and a child *v* is labeled with an actual syntenic region number, then vertex *v* induces an O event.
2. If a vertex *u* and its child *v* have actual syntenic region numbers and those two syntenic region numbers are different, then an R event is induced on the edge between *u* and *v*.

The objective of the DTLOR maximum parsimony reconciliation (DTLOR MPR) problem is to map the vertices and edges of the gene tree onto the species tree and to identify a syntenic region number with each internal vertex in the gene tree, minimizing the total cost of the events. Note that this model implicitly assumes that duplications are tandem or proximal duplications and thus a duplication event by itself does not imply a change of syntenic region. A duplication that gives rise to a copy at a different syntenic region is modeled implicitly by a duplication and a rearrangement event. The model can be extended to permit other types of duplication events.

### 2.2 Notation

Let *S* and *G* denote a pair of undated species and gene trees, respectively. Throughout this and the next section, we assume that *S* and *G* are binary. In Section 4 we extend results to non-binary trees.

For a tree *T*, let root(*T*) be the root and Le(*T*) be the set of leaves or *tips*. For a non-root vertex *v* in the tree, *p*(*v*) is the parent of *v*. For a non-leaf vertex *v*, *v*_1_ and *v*_2_ denote its two two children. We assume that each tree *T* has an additional *handle* edge, namely an edge (*u,* root(*T*)). The handle of *S* is denoted *e*^*S*^ and the handle of *G* is denoted *e*^*G*^. For a vertex *v* of *T*, we let *T* (*v*) be the subtree of *T* rooted at *v*, including its own handle edge *e*_*v*_ from *p*(*v*) to *v*. An edge of a tree *T* is said to be a *leaf edge* if its terminus is a leaf and is said to be an *internal edge* otherwise.

### 2.3 The DTLOR MPR Problem

An instance of the DTLOR-MPR problem is a 10-tuple (*S, G, L, ϕ, γ,* **D**, **T**, **L**, **O**, **R**) where:

- *S* = (*V*_*S*_, *E*_*S*_) and *G* = (*V*_*G*_, *E*_*G*_) are binary species and gene trees, respectively;
- *L* is a finite set of syntenic regions which are represented by counting numbers;
- *ϕ* : Le(*G*) → Le(*S*) is a function that maps the leaves of *G* to the leaves of *S*;
- *γ* : Le(*G*) → *L* is a surjective function that maps the leaves of *G* to syntenic regions;
- Parameters **D**, **T**, **L**, **O**, **R** are positive costs for duplication, transfer, loss, origin, and rearrangement events, described in detail below.

A *reconciliation* in the DTLOR model comprises a pair of functions (Φ, Γ) that extend the functions *ϕ* and *γ*. Specifically, Φ : *V* (*G*) → *V*(*S*) ∪ {*N*} maps the vertices of *G* to the vertices of *S* or the special *N* location representing a species that is not in the species tree *S*. The constraints on Φ are as follows:

1. Φ(*g*) = *ϕ*(*g*) for each leaf *g* of *G*;
2. If *g* is an internal vertex of *G* and Φ(*g*) ≠ *N* then the children of *g*, denoted *g*_1_ and *g*_2_, have the properties that

a. Φ(*g*_1_) ≠ *N* and Φ(*g*_2_) ≠ *N*;
b. Neither Φ(*g*_1_) nor Φ(*g*_2_) is an ancestor of Φ(*g*); and
c. At least one of Φ(*g*_1_) or Φ(*g*_2_) is equal to or a descendant of Φ(*g*).

Note that unlike the DTL model, which requires every gene vertex to be mapped to a vertex in the species tree, the DTLOR model allows gene vertices to be mapped to the *N* location which is outside the sampled species tree.

The mapping Φ induces four types of events. For an internal gene tree vertex *g*, with children *g*_1_ and *g*_2_, and Φ(*g*) ≠ *N*, the events induced by Φ are as follows:

#### Speciation event

Vertex *g* induces a speciation event if one of Φ(*g*_1_) and Φ(*g*_2_) is in the left sub-tree and the other is in the right subtree of Φ(*g*).

#### Duplication event

Vertex *g* induces a duplication event if each of Φ(*g*_1_) and Φ(*g*_2_) is either equal to or a descendant of Φ(*g*) but does not satisfy the requirements for a speciation event.

#### Transfer event

Vertex *g* induces a transfer event if exactly one of Φ(*g*_1_) and Φ(*g*_2_) is either equal to or a descendant of Φ(*g*) and the other is neither an ancestor nor a descendant of Φ(*g*).

#### Loss events

Each non-root vertex *g* (including leaf vertices) may induce zero or more loss events as follows: If Φ(*p*(*g*)) ≠ *N* is ancestral to Φ(*g*), then each species vertex *s* on the path from Φ(*p*(*g*)) to Φ(*g*) induces a loss event, except for Φ(*g*) and also not Φ(*p*(*g*)) if *p*(*g*) induces a speciation event. For each loss induced by a vertex *s* on the path from Φ(*p*(*g*)) to Φ(*g*), we say that *g passes* through *s*.

If Φ(*g*) = *N* then *g* induces none of these four types of events.

The function Γ : *V*(*G*) → *L* ∪ {*} maps each vertex *g* in *G* to an element of *L* or the special syntenic region represented by * indicating that it is an unknown location occurring outside of the species tree. The constraints on Γ and its relationship to Φ are as follows:

1. For *g* ∈ Le(*G*), Γ(*g*) = *γ*(*g*);
2. Φ(*g*) = *N* iff Γ(*g*) = *;
3. If Γ(*g*) ≠ * and *g* has children *g*_1_ and *g*_2_ then Γ(*g*_1_) ≠ * and Γ(*g*_2_) ≠ *.

The function Γ induces events as follows (recall that *p*(*g*) denotes the parent of *g*):

#### Origin event

A non-root vertex *g* induces an origin event if Γ(*p*(*g*)) = * and Γ(*g*) ≠ *. The root vertex root(*G*) induces an origin event if Γ(root(*G*)) ≠ *.

#### Rearrangement event

A non-root vertex *g* induces a rearrangement event if Γ(*g*) ≠ *, Γ(*p*(*g*)) ≠ *, and Γ(*p*(*g*)) ≠ Γ(*g*).

The cost of a reconciliation is defined to be the sum of the number of duplication, transfer, loss, origin, and rearrangement events scaled by the the event costs **D**, **T**, **L**, **O**, and **R** respectively. Speciation events are assigned an implicit cost of zero because a gene is expected to diverge when the species that carries it diverges.

## 3 The DTLOR MPR Algorithm

When a gene vertex *g* induces an origin event, all of the genes in the subtree *G*(*g*) rooted at *g* must have actual syntenic regions (by rule 3 in the definition of Γ), and the genes in that subtree are mapped to species in *S* (by rule 2 in the definition of Γ), that is, Φ(*g′*) ∈ *V*_*S*_ and Γ(*g′*) ∈ *L* for all *g′* ∈ *G*(*g*). The mappings Φ and Γ are only related by the constraint that Φ(*g*) = *N* iff Γ(*g*) = *. Thus, if *g* induces an origin event, then the pair of functions Φ and Γ restricted to the domain *G*(*g*) are independent. Therefore, for an *origin subtree*, a subtree of *G* whose root induces an origin event, the process of finding an optimal species mapping Φ can be decoupled from the process of finding an optimal location mapping Γ. Further, by definition, vertices that induce origin events cannot be ancestrally related. Thus, in a reconciliation (Φ, Γ) where *g′, g′′* induce origin events, the species and syntenic location mappings restricted to the origin subtree *G*(*g′*) are independent of the mappings restricted to the origin subtree *G*(*g′′*).

For binary gene trees, we use a dynamic programming algorithm to compute the optimal cost of a species mapping of each subtree of the gene tree. Then, we use a second dynamic programming algorithm to compute the optimal cost for the syntenic region mapping for each subtree. Finally, a third algorithm combines these results to find an optimal solution to the DTLOR MPR problem. For non-binary gene trees, this decoupling is no longer possible and a different (and less efficient) algorithm is presented in Section 4.

### 3.1 Computing the Species Map

Next, we give an efficient algorithm for computing an optimal species mapping for each origin subtree *G*(*g*). The algorithm is similar to other DTL reconciliation algorithms [4], but the variant used here is useful in the extensions and generalizations in later sections.

For a species mapping Φ, or its restriction to an origin subtree of the gene tree, we say that a gene tree edge *e*_*g*_ is *placed* on species tree edge *e*_*s*_ if either Φ(*g*) = *s* or if the path from Φ(*p*(*g*)) to Φ(*g*) includes vertex *s*. As a special case, if *g* is the root of an origin subtree, then Φ(*p*(*g*)) = *N*. In this case, there is no path from Φ(*p*(*g*)) to Φ(*g*), so *e*_*g*_ is placed on *e*_*s*_ if and only if Φ(*g*) = *s*. If Φ(*g*) = *s* we say that *e*_*g*_ *terminates* on edge *e*_*s*_ and if Φ(*g*) is a descendant of *s* then a loss event is induced and we say that *e*_*g*_ *continues* on the corresponding child edge of *e*_*s*_.

Let *C*(*g*) denote the optimal cost for a species mapping restricted to the domain of *G*(*g*) and let *C*(*e*_*g*_, *e*_*s*_) denote the optimal cost for a species mapping of *G*(*g*) such that *e*_*g*_ is placed on *e*_*s*_. Then 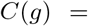 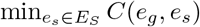.

We now describe an algorithm for computing *C*(*e*_*g*_, *e*_*s*_). The algorithm computes the *C* table by considering edges in the gene tree bottom-up (postorder): An edge *e*_*g*_ is considered if either *g* is a leaf or the children edges 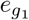 and 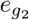 have already been considered. For each edge *e*_*g*_ under consideration, we now consider each edge *e*_*s*_ in the species tree in postorder.

To compute *C*(*e*_*g*_, *e*_*s*_), we enumerate the four possible cases:

- In the base case, if *g* and *s* are leaves, then:

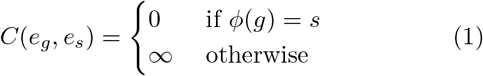
- If neither *g* nor *s* is a leaf, then either *g* is mapped to *s* or not. If *g* is not mapped to *s*, then it induces a loss at *s* by being mapped to one of its children. Otherwise, *g* is mapped to *s*, which induces either a cospeciation, duplication, or transfer event, which incurs a corresponding cost. Thus,

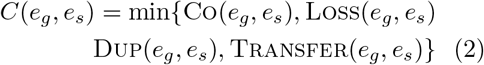

where the computation of Co, Loss, Dup, and Transfer are described below.
- If *g* is not a leaf but *s* is a leaf, then cospeciation and loss at *g* are not possible, so:

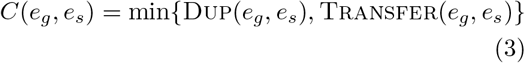
- If *g* is a leaf but *s* is not a leaf, then cospeciation, duplication and transfer at *g* are not possible, so:

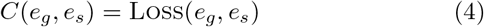

The functions Co(*e*_*g*_, *e*_*s*_), Loss(*e*_*g*_, *e*_*s*_), Dup(*e*_*g*_, *e*_*s*_), and Transfer(*e*_*g*_, *e*_*s*_) are computed as follows:

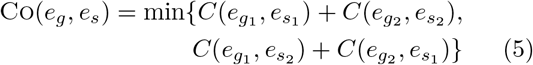

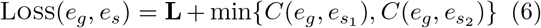

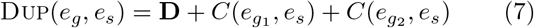

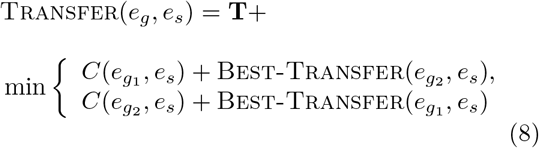

The cospeciation term (5) considers both ways in which the children edges of *e*_*g*_ can be placed onto the children edges of *e*_*s*_ in a cospeciation event. The loss term (6) considers both ways in which edge *e*_*g*_ can continue, either on one child of *e*_*s*_ or the other. The duplication term (7) places both children edges of *e*_*g*_ on *e*_*s*_. In the transfer term (8), we consider both ways of selecting the transferred child edge. The non-transferred child edge of *e*_*g*_ remains on *e*_*s*_, but the transferred child edge is placed on a species edge determined by Best-Transfer; Best-Transfer(*e*_*j*_, *e*_*s*_) denotes the minimum cost of a mapping of the subtree *G*(*g*_*j*_) assuming that *e*_*j*_ is placed on a species edge that is neither ancestral nor descendant to *e*_*s*_. In order to compute these values, the algorithm maintains another table called Best-Entry(*e*_*g*_, *e*_*s*_) which stores the minimum of *C*(*e*_*g*_, *e*_*i*_) over all *e*_*i*_ in the subtree rooted at *e*_*s*_. The algorithm is given in Algorithm 1.

**Algorithm 1:**
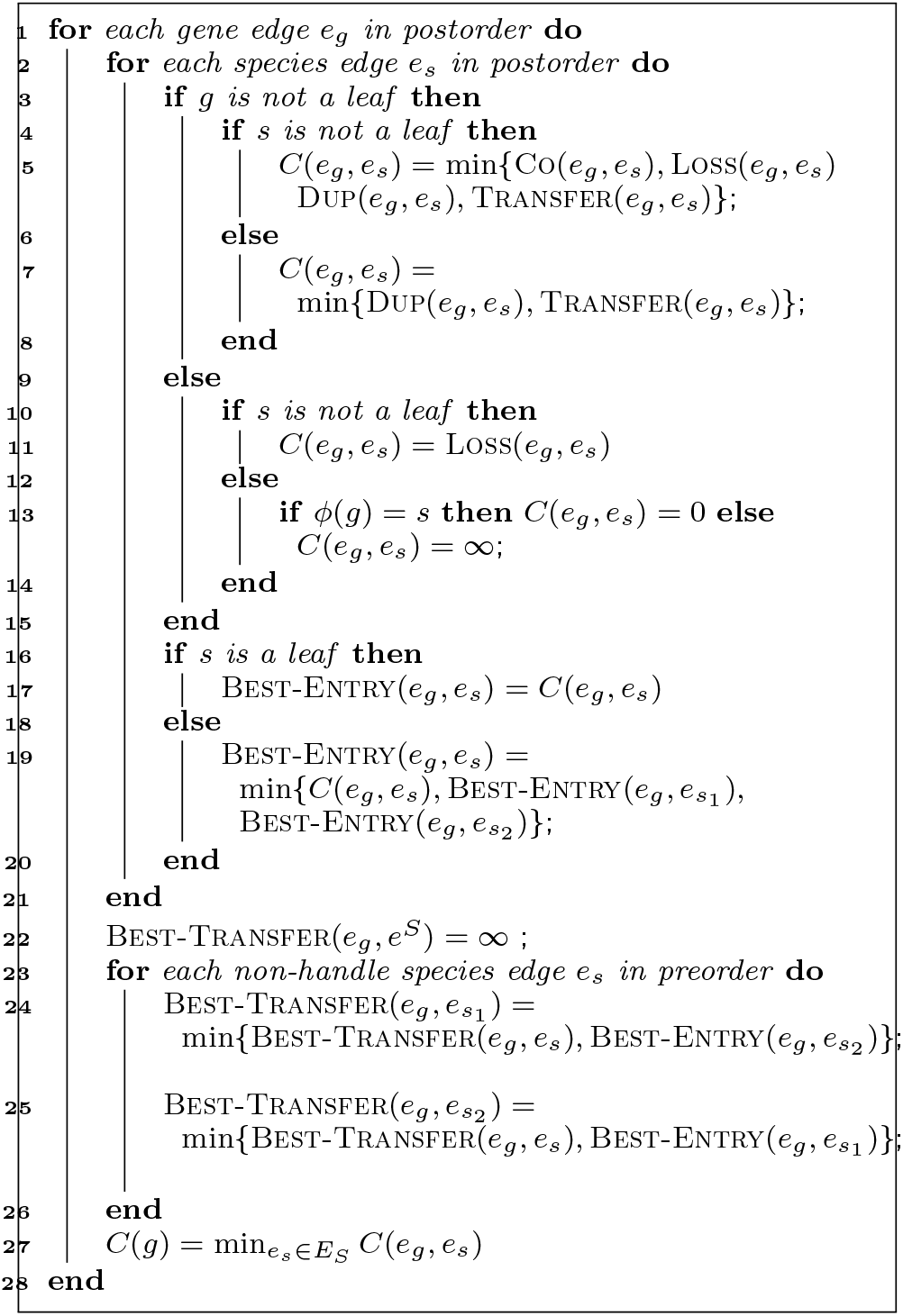
Computing *C*(*g*)

### 3.2 Computing the Synteny Map

We use another dynamic programming algorithm to find the optimal cost for a syntenic region mapping for each subtree *G*(*g*). Let syn(*g*) denote the optimal cost for a syntenic region mappping of *G*(*g*). Let syn(*g, ℓ*) denote the optimal cost for a syntenic region mapping of *G*(*g*) such that *g* has the syntenic region *ℓ*. Then syn(*g*) = min_*ℓ*∈*L*_ syn(*g, ℓ*).

If *g* is a leaf, then Γ(*g*) = *γ*(*g*) (by rule 1 in the definition of Γ). Thus,

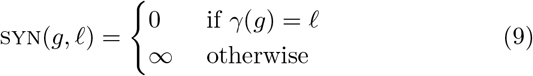

If *g* is not a leaf, then (recalling that *g*_1_ and *g*_2_ denote the children of *g*):

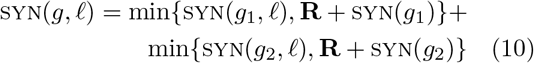

This accounts for each child *g*_1_ and *g*_2_ either remaining in the same syntenic region as *g* or potentially changing to a new region and incurring a cost of **R**. The algorithm for computing syn(*g, ℓ*) is summarized in Algorithm 2.

**Algorithm 2:**
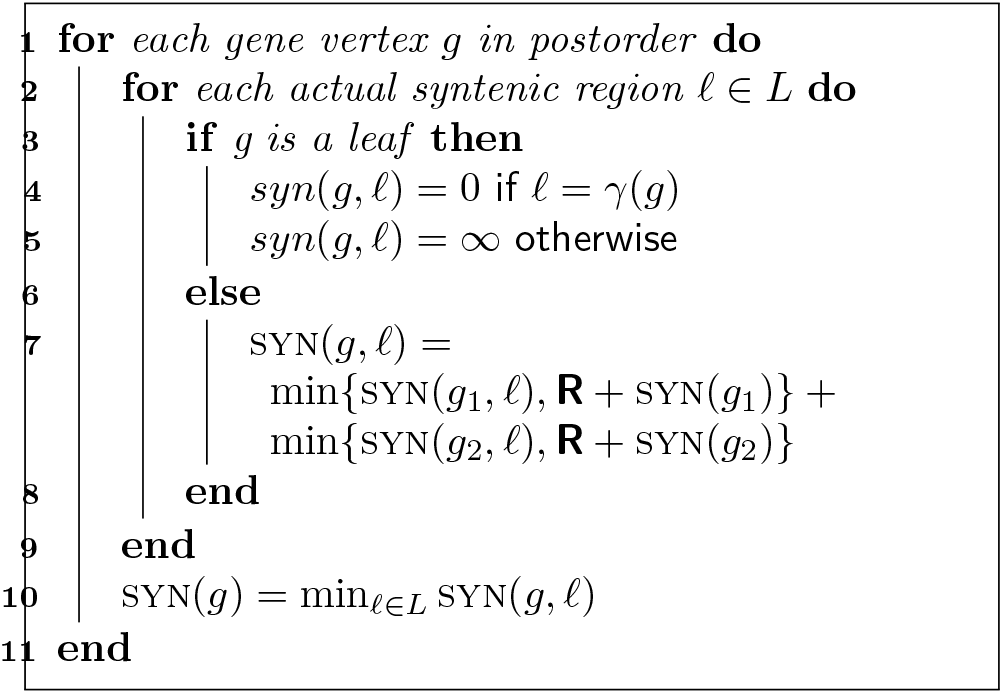
Computing syn(*g*)

### 3.3 Solving the DTLOR MPR Problem

Let Origin(*g*) denote the cost of reconciling an origin subtree *G*(*g*) which, as noted earlier, can be computed as Origin(*g*) = **O**+*C*(*g*)+syn(*g*). To find a maximum parsimony reconciliation, we must therefore determine the optimal locations for origin events.

Let Null(*g*) be the optimal cost of reconciling *G*(*g*) such that *g* has the unknown syntenic region *. Since *γ* must be respected, *g* may not be assigned syntenic region * if *g* is a leaf. Thus, Null(*g*) is calculated as:

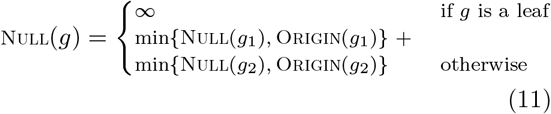

The optimal cost for reconciling the entire gene tree *G* is given by:

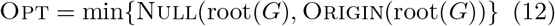

The algorithm for computing Null(*g*) is summarized in Algorithm 3.

**Algorithm 3:**
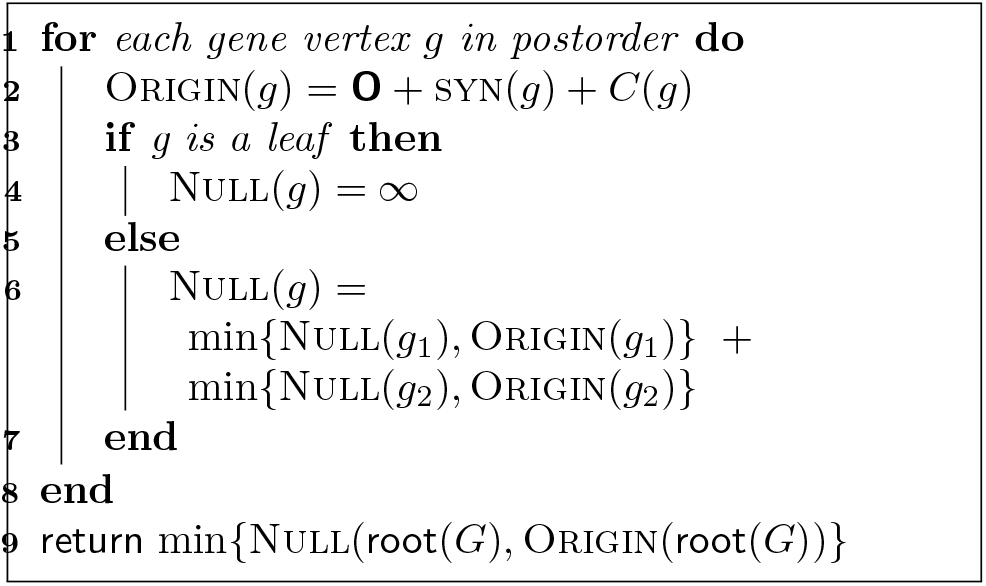
Computing the DTLOR MPR Solution

Note that if we wish to reconstruct an optimal solution, the DP tables *C,* syn, Null can be annotated in the standard way, allowing the solutions to be reconstructed by tracing through the table. We first trace through the Null table to find a set of origin events that produce an optimal solution. For any gene vertex not in any of the origin subtrees induced by these origin events, they are labeled with the unknown syntenic region *. For each origin subtree, we trace through the syn table to get an optimal location mapping, and then we trace through the *C* table to get an optimal species mapping. Because of loss events, there can be multiple *C*(*e*_*g*_, *e*_*s*_) entries that involve the same gene vertex in an optimal solution. The mapping of that gene vertex corresponds to the lowest such *e*_*s*_.

The proofs of the following are given in the Appendix:

#### Lemma 1

*Algorithm 1 correctly computes C*(*g*) *for every gene vertex g* ∈ *V* (*G*).

#### Lemma 2

*Algorithm 2 correctly computes* syn(*g*) *for every gene vertex g* ∈ *V* (*G*).

#### Theorem 1

*Algorithm 3 correctly computes the optimal solution to the DTLOR-MPR problem.*

### 3.4 Time Complexity

Computing each entry of the C table takes constant time, so the running time for computing the C table is *O*(|*G||S*|). Computing the syn table takes *O*(|*G||L*|) time, and computing the Origin and Null entries takes *O*(|*G*|) time. In total, the asymptotic running time of this algorithm is *O*(|*G*||*S*| + |*G*||*L*|).

## 4 Non-binary Gene Trees

While it is generally possible to construct accurate species trees using a variety of methods, gene trees are susceptible to ambiguity due to the relatively little information available in their sequence data. Consequently, phylogenetic trees for genes often have non-binary vertices, also known as *multifurcations* or *soft polytomies*, which correspond to an unknown ordering of the underlying sequence of divergences [11]. In this case, we wish to expand each multifurcation into a sequence of binary divergences, leading to a binary gene tree. Such an expansion is called a *resolution* or *binarization* of the non-binary tree. Our goal is to find a resolution of the gene tree that leads to a lowest cost solution to the DTLOR MPR problem.

Unfortunately, the number of resolutions of a non-binary tree can be exponential in the number of vertices in the tree. It is, therefore, impractical to explicitly consider every resolution. However, Kordi and Bansal [11] and Jacox *et al.* [12] demonstrated the existence of fixed-parameter polynomial-time algorithms for finding maximum parsimony reconciliations for non-binary trees in the DTL model. These algorithms operate in polynomial time assuming the maximum number of children of any non-binary vertex is bounded by some constant *k*. More precisely, a fixed-parameter algorithm in this context runs in time *O*(*f* (*k*)*p*(*m, n*)) where *m* and *n* denote the sizes of the gene and species trees, *k* is the maximum branching factor of any gene vertex, *p*(*m, n*) is a polynomial in *m* and *n*, and *f* (*k*) is some function of *k* which may even be exponential in *k*. In particular, *f* (*k*) in this context is the number of distinct binary resolutions of a tree comprising a root and *k* children which can be shown to be *f*(*k*) = 2^*k*^(*k* − 1)!. For any fixed *k*, this value is a fixed constant. Jacox *et al.* [12] offer an approach that results in an *f* (*k*) which is smaller but still potentially exponential in *k*. Importantly, a fixed-parameter polynomial-time algorithm is much more efficient and practical than an exponential time algorithm such as the naive approach of enumerating all possible resolutions of a non-binary tree which would have running time *O*((2^*k*^(*k* − 1)!)^*n*^).

In this section, we describe a fixed-parameter polynomial time algorithm for the DTLOR MPR problem. Following the approach of Kordi and Bansal [11], our algorithm expands each individual non-binary vertex into every possible binary resolution but avoids enumerating all possible binary resolutions of the entire tree, thus resulting in a fixed-parameter polynomial-time algorithm rather than an exponential-time algorithm. While our algorithm leverages the important ideas for resolving non-binary vertices first proposed by Kordi and Bansal [11], it requires a new algorithm due to the advent of *O* and *R* events.

### 4.1 Description of the Algorithm

Each binary resolution of a non-binary gene tree implies a different topology for the gene tree, which induces potentially different costs for both the species mapping and the syntenic region mapping. In particular, one resolution may be most favorable for minimizing the cost of the species mapping while a different one may admit the least expensive syntenic region mapping (Figure 1). Thus, while in binary gene trees the species mapping and syntenic region mappings could be efficiently solved independently and then merged into an optimal solution for the DTLOR MPR problem, the situation is more complicated in the presence of non-binary gene trees. The algorithm presented here considers the species mapping and syntenic region mapping simultaneously as non-binary vertices are resolved one-by-one.

**Figure 1.**
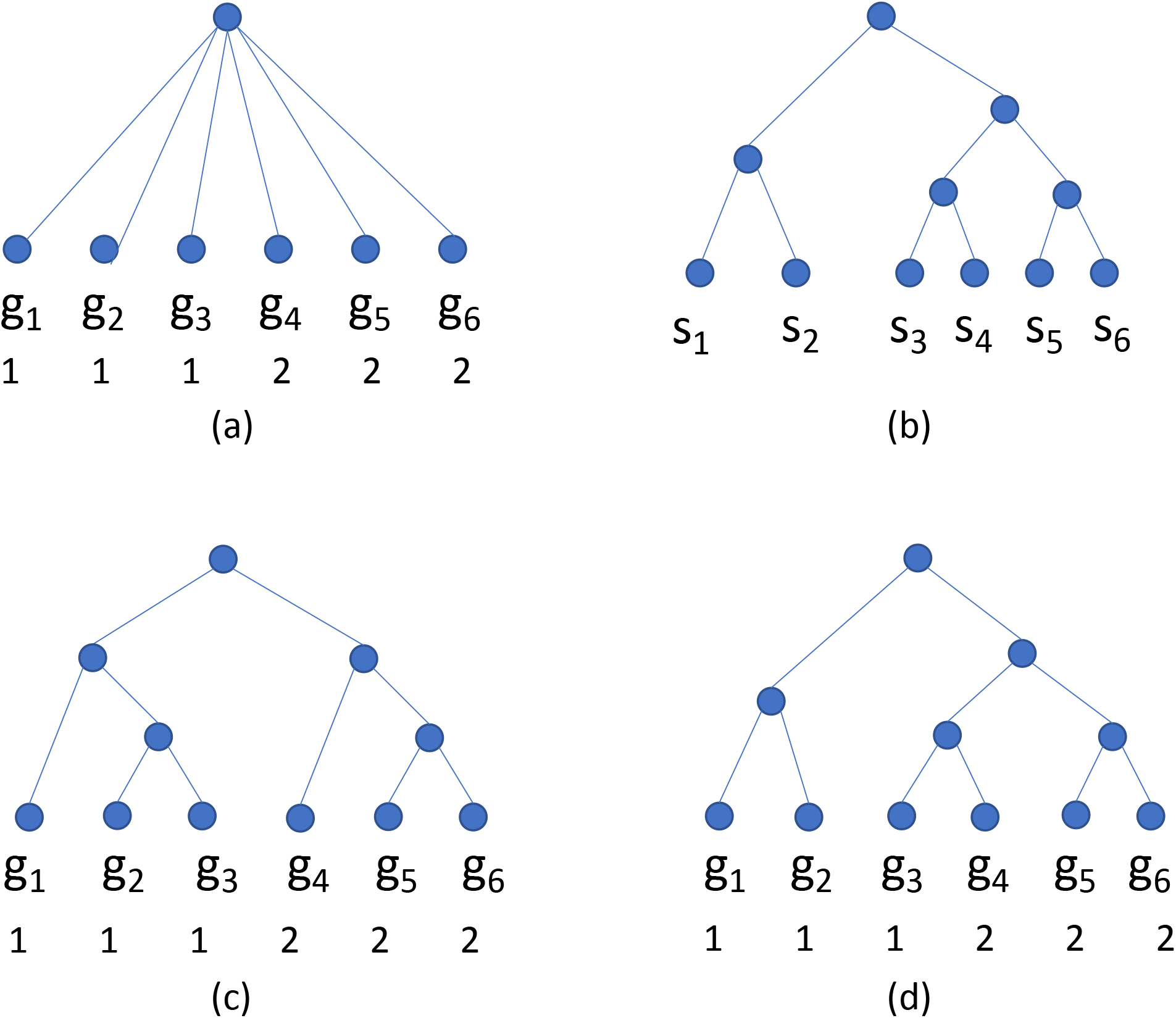
An example showing that one resolution of a multifurcation can be optimal for species mapping while another may be optimal for syntenic region mapping. (a) A gene tree with six leaves labeled with their syntenic regions. (b) A species tree. The tip association is *ϕ*(*g*_*i*_) = *s*_*i*_, 1 ≤ *i* ≤ 6. (c) A binary resolution of the gene tree in which the optimal species mapping cost is necessarily greater than the Origin cost **O** since this tree is not isomorphic to the species tree; the optimal number of rearrangements in this case is 1. (d) Another binary resolution of the gene tree in which the optimal species mapping cost is just the Origin cost **O** since this tree is isomorphic to the species tree; the optimal number of rearrangements in this case is 2.

Let *C*(*e*_*g*_, *e*_*s*_, *ℓ*) denote the optimal cost of reconciling the subtree *G*(*g*) with *S* such that *e*_*g*_ is placed on *e*_*s*_ and *g* has the syntenic region *ℓ*. Note that in contrast to *C*(*e*_*g*_, *e*_*s*_) used in the previous section, *C*(*e*_*g*_, *e*_*s*_, *ℓ*) also encodes the constraint that *g* has syntenic region *ℓ* and the total cost includes the cost of rearrangement events in the subtree *G*(*g*). We define *O*(*e*_*g*_, *e*_*s*_, *ℓ*) and Best-Transfer(*e*_*g*_, *e*_*s*_, *ℓ*) analogously to *O*(*e*_*g*_, *e*_*s*_) and Best-Transfer(*e*_*g*_, *e*_*s*_), respectively, in the previous section. We define *C*(*e*_*g*_, *e*_*s*_, *L*) = min_*ℓ*∈*L*_ *C*(*e*_*g*_, *e*_*s*_, *ℓ*).

We compute *C*(*e*_*g*_, *e*_*s*_, *ℓ*) in postorder. There are four cases:

- In the base case, if *g* and *s* are leaves, then:

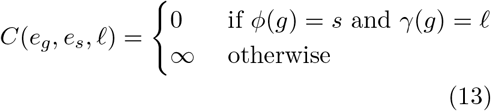
- If neither *g* nor *s* is a leaf, then:

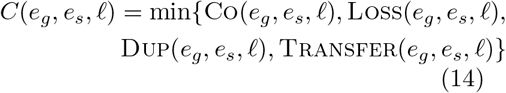

where the computation of Co, Loss, Dup, and Transfer are given below.
- If *g* is not a leaf but *s* is a leaf, then:

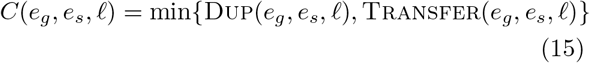
- If *g* is a leaf but *s* is not a leaf, then:

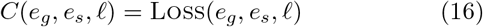

The functions Co(*e*_*g*_, *e*_*s*_, *ℓ*), Loss(*e*_*g*_, *e*_*s*_, *ℓ*), Dup(*e*_*g*_, *e*_*s*_, *ℓ*), and Transfer(*e*_*g*_, *e*_*s*_, *ℓ*) are computed as follows:

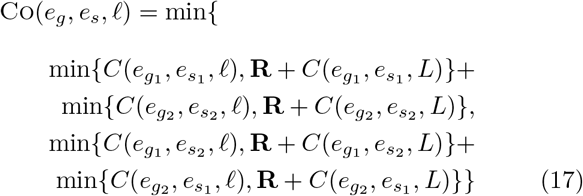

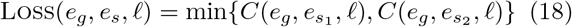

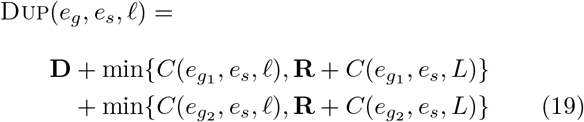

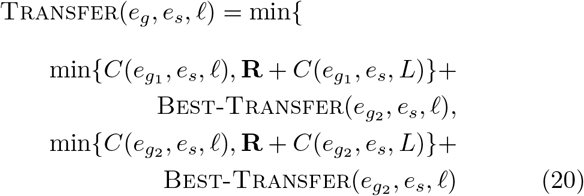

In order to compute Best-Transfer, we compute Best-Entry(*e*_*g*_, *e*_*s*_, *ℓ*) as follows. If *s* is a leaf, then

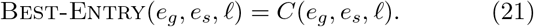

Otherwise,

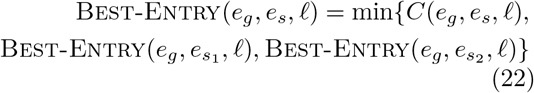

Best-Transfer(*e*_*g*_, *e*_*s*_, *ℓ*) is then computed in pre-order: First, for the handle edge *e*^*S*^

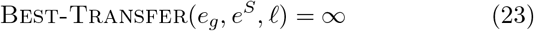

For all other edges, *e*_*s*_ with child branches 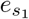 and 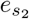

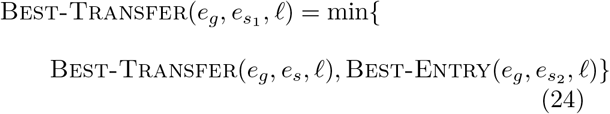

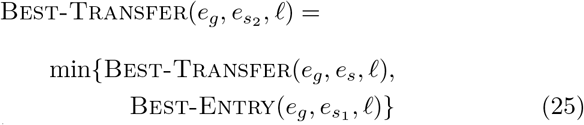

For a gene vertex *g* with children *g*_1_*, g*_2_*, . . ., g*_*k*_, *k >* 2, a binary resolution for *g* is defined to be a binary tree whose root is *g* and whose leaves are *g*_1_*, g*_2_*, . . ., g*_*k*_. Let *BR*(*g*) denote the set of all binary resolutions for *g*. Let Null(*g*) be the optimal cost of reconciling *G*(*g*) such that *g* has the unknown syntenic region *. Let Origin(*g*) be the optimal cost of reconciling *G*(*g*) such that *g* induces an origin event. Let *A*^*H*^, *C*^*H*^, Best-Entry^*H*^, Best-Transfer^*H*^, Origin^*H*^, Null^*H*^ correspond to *A*,*C*, Null for a specific binary resolution *H* for *g*. Let *e*_*h*_ be an edge in *H*. If *e*_*h*_ is a leaf edge in *H* (and thus *h* is one of the children of *g* in *G*), then *C*^*H*^(*e*_*h*_, *e*_*s*_, *ℓ*) = *C*(*e*_*h*_, *e*_*s*_, *ℓ*) for all *e*_*s*_, *ℓ*, Origin^*H*^(*h*) = Origin(*h*), and Null^*H*^(*h*) = Null(*h*).

The algorithm for reconciling a non-binary gene tree *G* with a species tree *S* is summarized in Algorithm 4. All DP table entries are initialized to ∞.

#### Theorem 2

*Algorithm 4 correctly computes the optimal solution to the DTLOR-MPR problem for non-binary gene trees.*

The proof is analogous to the proof of correctness of Theorem 1.

### 4.2 Time Complexity

The algorithm first initializes Origin, Null, *C* entries for all the leaf edges of the gene tree, which takes *O*(|*G*|). Then for each internal gene edge *e*_*g*_, it loops through all binary resolution *H* at the gene vertex and computes *A*^*H*^, *C*^*H*^, Best-Entry^*H*^, Best-Transfer^*H*^, Origin^*H*^, and Null^*H*^, which takes *O*(|*H||S||L|*) = *O*(*k|S||L|*) time. Here *|H|* is the size of any binary resolution at a gene vertex, which is bounded by *O*(*k*) where *k* is the maximum degree of the gene tree. For each *H*, it also updates all entries of *C*(*e*_*g*_, *e*_*s*_, *ℓ*) and *C*(*e*_*g*_, *e*_*s*_, *L*), which takes *O*(|*L||S|*) time. In total, the running time for computing all the DP entries for all gene edges and binary resolutions is *O*(*f*(*k*)*k|G|S||L|*) time, where *f*(*k*) upper bounds the number of binary resolutions at any gene vertex.

## 5 Integration in xenoGI

The implementation of the DTLOR MPR Algorithm (Algorithm 3) was integrated into the xenoGI software package [2] which seeks to reconstruct the history of genome evolution in clades of microbes. xenoGI takes as input a set of sequenced genomes, identifies gene families within this set, and groups those families by common origin. The previous version of xenoGI was able to map gene families onto the species tree and identify their point of origin. However, it was not able reconstruct events in the subsequent evolution of the gene family (e.g.losses or rearrangements). The integration of the DTLOR MPR algorithm allows xenoGI to reconstruct these subsequent events, providing potentially important new insights into microbial evolution.

**Algorithm 4:**
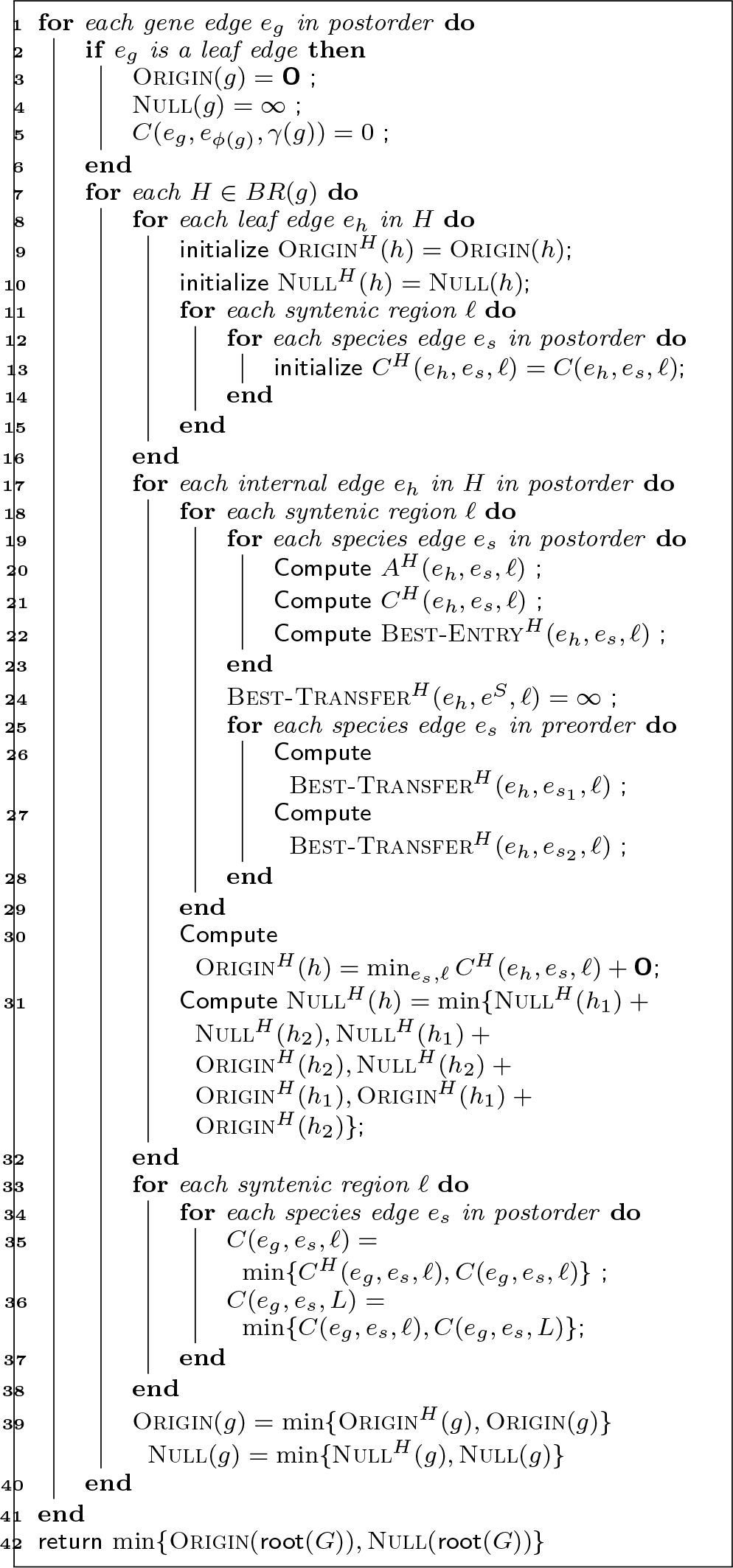
The DP algorithm for non-binary gene trees.

Within the new DTLOR version of xenoGI, we construct a gene tree for every family, then reconcile it with the species tree. The resulting reconciliation can be used to refine the family (e.g. split it into multiple parts based on the placement of origin events) and to provide detailed information about the family’s subsequent evolution.

The table below shows the running time for the DTLOR MPR Algorithm within xenoGI on all gene trees given inputs ranging from 4 to 15 bacterial genomes (species). In each case, DTLOR was run on every binary gene tree with more than two leaves. These calculations were performed on a commodity server (50 AMD Opteron 6276 2.3 GHz processors, 503 GB RAM). The DTLOR costs were set at 1, 1, 1, 2, 2, respectively.

In one of our enteric bacterial test data sets (Dataset B in Table 1) we examined DTLOR output for a known genomic island, the Acid Fitness Island (AFI) [13]. The corresponding species tree is shown in Figure 2. This island appears to have originated with an insertion of 19 genes on branch s2 (the branch leading to the internal node s2) via horizontal transfer from outside the clade [2]. It then evolved in the clade and was inherited in the the four descendant strains, with the notable loss of nine genes along the branch leading to E. coli K12. For nearly all the gene families in this island, DTLOR produced reconciliations that place the origin of the family on the branch leading to s2, and it correctly recognized loss events on the K12 branch where those occurred. In a few cases, there were multiple most parsimonious reconciliations (MPRs), one of which was deemed correct, and the others incorrect. Finally there is one family (glutamate decarboxylase) whose complicated post-insertion evolution is hard even for a human expert to sort out. In this case none of the MPRs appear correct. (The evolution of this family likely involved gene conversion, but the MPR lacks T events using the event costs that we used in this experiment.)

**Table 1.**
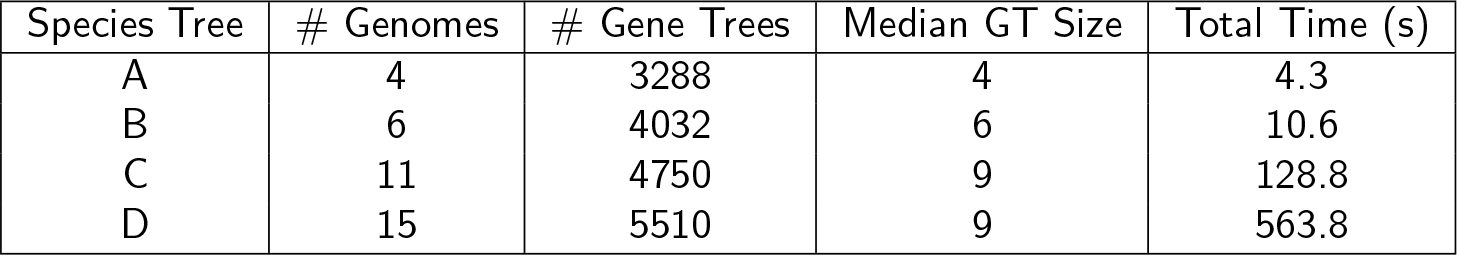
Runtimes for four different species trees with 4, 6, 11 and 15 tips.

**Figure 2.**
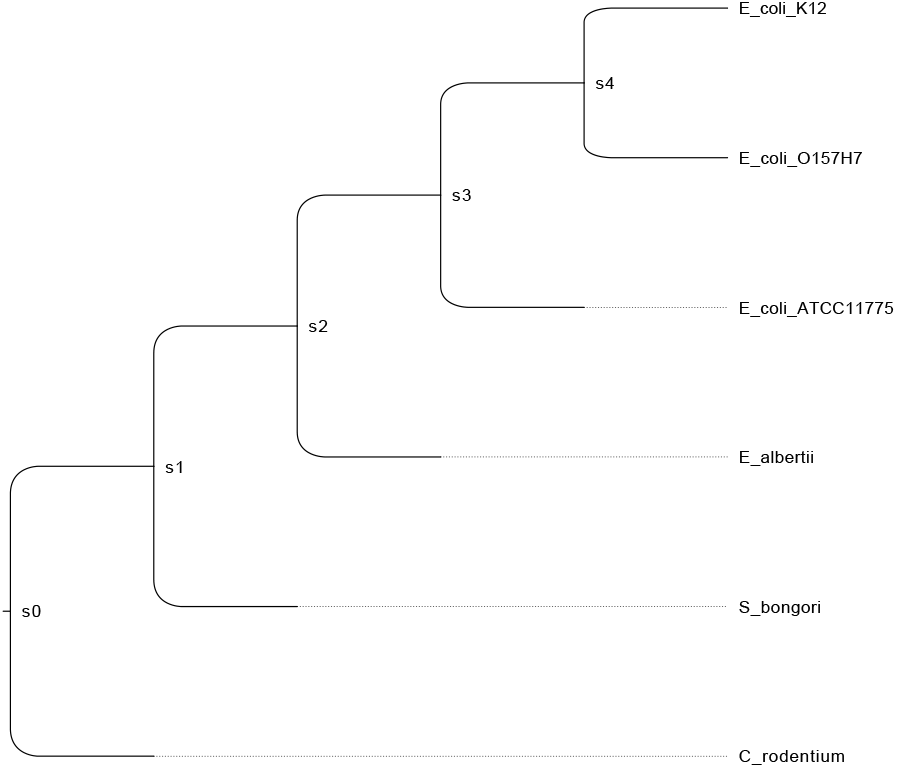
The species tree for Dataset B.

## 6 Conclusion and Future Work

In this paper, we have described the DTLOR event model which extends the well-known DTL model to include origin and rearrangement events. This model is particularly applicable to the evolution of microbes where the species tree is, in many cases, not fully sampled. Therefore, reconciliations must be able to account for transfer events from outside the sampled tree. In addition, the DTLOR model allows for syntenic rearrangement, which is also prevalent in microbial gene families.

We have described efficient algorithms for maximum parsimony reconciliation in the DTLOR model. In binary gene trees, our algorithm solves the DTL reconciliation problem and the sytnenic region problems independently and then combines the results of those two algorithms, resulting in a particularly efficient solution. When the gene tree is non-binary, the two sub-problems can evidently no longer be decoupled in this way, and our algorithm for this case considers all of the events simultaneously.

Several important problems remain to be studied. First, the impact of event costs is not well-understood. Just as in the case of the DTL model, different event costs can give rise to different reconciliations which, in turn, can lead to different conclusions. We believe that the costscape algorithm developed for the DTL model [14] may be extendible to the DTLOR model, which would provide insights into the impact of event costs to the solution space.

Second, it is often the case that there are many distinct MPRs. In fact, even in the DTL model, the number of MPRs can be exponential in the size of the two trees [15]. A subset of these MPRs may be of particular interest because they contain certain evolutionary events that are strongly believed to have occurred (e.g., a horizontal transfer on a particular branch of the species tree). It is desirable, therefore, to filter the set of MPRs to only include those that contain a specified set of events, count the number of MPRs in the filtered set, compute support values on the constituent events in that space of MPRs, and select representative reconciliations from this set.

Finally, further systematic studies are needed to determine the full impact of the DTLOR MPR algorithm on the analyses that can now be performed with the enhanced xenoGI tool, including the case of non-binary gene trees.

## Abbreviations

DTLOR: Duplication-Transfer-Loss-Origin-Rearrangement
MPR: Maximum Parsimony Reconciliation

## Declarations

### Ethics approval and consent to participate

Not applicable.

### Consent for publication

Not applicable.

### Availability of data and materials

The DTLOR algorithm for binary trees has been implemented in Python 3 and is available for download at https://github.com/aremath/DTLOR.

## Authors’ contributions

RLH and EB conceived this research. SS made contributions to the model. RLH and NL performed the first implementation of the algorithm. JL and RM made contributions to the development and analysis of the algorithms, analysis, and proofs of correctness. JL, RM, EB, and RLH wrote the paper.

## Acknowledgements

The authors thank Yi-Chieh Wu for valuable discussions.

## Author details

^1^ Department of Computer Science, Harvey Mudd College, Claremont, California, USA. ^2^ Department of Biology, Harvey Mudd College, Claremont, California, USA. ^3^ Department of Computer Science and Engineering, University of Califorina Santa Cruz, Santa Cruz, California, USA.

## Appendix

This appendix contains proofs of results from the main paper.

### Proof of Lemma 1

*Proof* We prove that the algorithm correctly computes *C*(*e*_*g*_, ·) and Best-Transfer(*e*_*g*_, ·) by structural induction on *G*. For the base case for *C*(*e*_*g*_, ·), consider a leaf edge *e*_*g*_. We perform structural induction on *S*. In the base case where *e*_*s*_ is a leaf edge, *g* must map to *ϕ*(*g*), so *C*(*e*_*g*_, *e*_*s*_) is computed correctly by equation 1. In the inductive step, consider a non-leaf edge *e*_*s*_. In this case, *g* must induce a loss event at *s* since *g* is a leaf, so *C*(*e*_*g*_, *e*_*s*_) = Loss(*e*_*g*_, *e*_*s*_) (by equation 2). By the inductive hypothesis, for each descendant branch *e*_*s′*_ of *e*_*s*_, *C*(*e*_*g*_, *e*_*s′*_) is computed correctly, so Loss(*e*_*g*_, *e*_*s*_) is computed correctly (by equation 6). This concludes the base case for *C*(*e*_*g*_, ·).

In the base case for Best-Transfer(*e*_*g*_, ·), we consider a leaf edge *e*_*g*_. Since its correctness relies on Best-Entry(*e*_*g*_, ·), we first use structural induction on *S* to prove that Best-Entry(*e*_*g*_, ·) is correctly computed. In the base case, *e*_*s*_ is a leaf edge, so the only choice for *e*_*g*_ to enter the subtree rooted at *e*_*s*_ is at *e*_*s*_. Since *C*(*e*_*g*_, *e*_*s*_) is correctly computed, Best-Entry(*e*_*g*_, *e*_*s*_) is also correctly computed on line 17. In the inductive step, consider a non-leaf edge *e*_*s*_. By the inductive hypothesis, Best-Entry(*e*_*g*_, *e*_*s′*_) is computed correctly for each descendant edge *e*_*s′*_ of *e*_*s*_. The ways for *e*_*g*_ to enter the subtree rooted at *e*_*s*_ are at *es*, the left subtree of *es*, or the right subtree of *e*_*s*_, thus Best-Entry(*e*_*g*_, *e*_*s*_) is computed correctly on line 19.

Now we prove the base case for Best-Transfer(*e*_*g*_, ·) using structural induction on *S* from the handle edge *e*^*S*^. In the base case where *e*_*s*_ = *e*^*S*^, all edges in *S* is a descendent of *e*^*S*^, so there is no valid species edge for the child edge of *e*_*g*_ to transfer to. Thus Best-Transfer(*e*_*g*_, *e*^*S*^) is computed correctly on line 22. In the inductive step, we consider a non-root edge 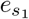, which has a sibling edge 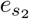. By the inductive hypothesis 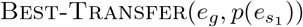 is computed correctly and by the inductive proof on Best-Entry(*e*_*g*_, ·), 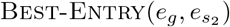 is computed correctly. Since the species edges that *e*_*g*_ are allowed to transfer to from 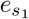 not only include the same edges if *e*_*g*_ were to transfer from 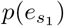, but also edges in the subtree rooted at 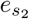, and the optimal cost of placing *e*_*g*_ inside the subtree rooted at 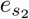 is given by 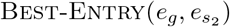, Best-Transfer(*e*_*g*_, *e*_*s*_) is computed correctly. This concludes the base case for Best-Transfer(*e*_*g*_, ·). In the inductive step, consider a non-leaf edge *e*_*g*_. By the inductive hypothesis *C*(*e*_*g′*_, ·) and Best-Transfer(*e*_*g′*_, ·) are computed correctly for every descendant edge of *e*_*g′*_ of *e*_*g*_. Thus, Best-Transfer(*e*_*g*_, ·) is computed correctly on lines 24 and 25.

To conclude the proof of correctness of *C*(*e*_*g*_, ·), we use structural induction on *S*. In the base case, *e*_*s*_ is a leaf edge; the only two possibilities are *e*_*g*_ duplicates on *e*_*s*_ or transfers on *e*_*s*_. The correctness of Dup(*e*_*g*_, *e*_*s*_) is guaranteed by the the correctness of *C*(*e*_*g′*_, ·), while the correctness of Transfer(*e*_*g*_, *e*_*s*_) is guaranteed by the correctness of both *C*(*e*_*g′*_, ·) and Best-Transfer(*e*_*g′*_, ·) for each descendant edge *e*_*g′*_ of *e*_*g*_. In the inductive step, *e*_*s*_ is not a leaf edge, then *C*(*e*_*g*_, *e*_*s*_) is, by definition, the minimum of Co(*e*_*g*_, *e*_*s*_), Dup(*e*_*g*_, *e*_*s*_), Transfer(*e*_*g*_, *e*_*s*_), and Loss(*e*_*g*_, *e*_*s*_) (see equation 4). Again, Co(*e*_*g*_, *e*_*s*_), Dup(*e*_*g*_, *e*_*s*_) and Transfer(*e*_*g*_, *e*_*s*_) are computed correctly by the correctness of *C*(*e*_*g′*_, ·) and Best-Transfer(*e*_*g′*_, ·). The correctness of Loss(*e*_*g*_, *e*_*s*_) is guaranteed by the inductive hypothesis on the correctness of *C*(*e*_*g*_, *e*_*s′*_) for every descendant edge *e*_*s′*_ of *e*_*s*_. This concludes the inductive step for *C*(*e*_*g*_, *e*_*s*_).

Finally, *C*(*g*) is the optimal cost for a species mapping of *G*(*g*) such that *g* can be mapped to any species in *S*. By definition, 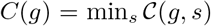 where 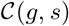 denotes the optimal cost for a species mapping of *G*(*g*) in which *g* is mapped to *s*.

Consider 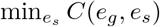. Since *C*(*e*_*g*_, *e*_*s*_) is the optimal cost of a mapping where *e*_*g*_ is placed on *e*_*s*_, this implies a mapping of *g* to *s* or one of its descendants. If *g* is mapped to *s*, then 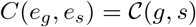. If *C*(*e*_*g*_, *e*_*s*_) involves a mapping of *g* to *s′* which is a descendant of *s*, then it induces loss events. Because loss events have a non-negative cost, *C*(*e*_*g*_, *e′*_*s*_) ≤ *C*(*e*_*g*_, *e*_*s*_), and thus 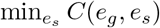 only includes those entries of *C*(*e*_*g*_, *e*_*s*_) where *g* is mapped to *s*. Thus, 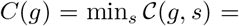 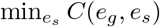.

### Proof of Lemma 2

*Proof* We prove the correctness of syn(*g, ℓ*) and syn(*g*) by induction on *g*. In the base case, *g* is a leaf, then the syntenic region of *g* must be *γ*(*g*), so both syn(*g, ℓ*) and syn(*g*) are computed correctly on line 5. In the inductive step, consider an internal vertex *g*. By the inductive hypothesis, syn(*g*_*i*_, *ℓ′*) and syn(*g*_*i*_) is computed correctly for any *g*_*i*_ ∈ *Ch*(*g*) and *ℓ′* ∈ *L*. The syntenic region for the left and the right child are chosen independently, so the total cost for syn(*g, ℓ*) is the sum of the costs for choosing a location for each child of *g*. If *g*_*i*_ is a child of *g* with syntenic region *ℓ*, then no *R* event is induced and the cost is syn(*g*_*i*_, *ℓ*). If an *R* event is induced by a change in syntenic region, then it is optimal to choose a location that minimizes the total cost of choosing syntenic regions for *G*(*g*_*i*_). Thus the cost for *g*_*i*_ with an *R* event is syn(*g*_*i*_) + **R** The cost for *g*_*i*_ as whole is the minimum of these two possibilities, and this is the same for both children of *g*. Thus, syn(*g, ℓ*) is computed correctly on line 7. Then, by definition, syn(*g*) is also computed correctly by taking the minimum of syn(*g, ℓ*) over all *ℓ* ∈ *L*.

### Proof of Theorem 1

*Proof* First we prove the correctness of Origin(*g*). Since a species mapping for an origin subtree *G*(*g*) is independent of a location mapping of the same sub-tree, the optimal cost of a reconciliation of *G*(*g*) is the optimal cost *C*(*g*) for a species mapping and the optimal cost syn(*g*) for a location mapping and the cost of the Origin event at *g*. Thus Origin(*g*) is computed correctly on line 2.

Now we prove Null(*g*) is computed correctly by an induction on *g*. In the base case, *g* is a leaf, so *g* must have a known location *γ*(*g*). Thus Null(*g*) = ∞ as computed on line 4. In the inductive case, consider an internal vertex *g*. By the inductive hypothesis, Null(*g*_*i*_) is computed correctly for any *g*_*i*_ ∈ *Ch*(*g*). There are four cases we need to consider since each child can induce an Origin event or remain unassigned to any real syntenic region. Since we already know Origin(*g*_*i*_) and Null(*g*_*i*_) are computed correctly, taking the minimum over the four cases yield the optimal cost for assigning *g* to the unknown syntenic region *. Thus Null(*g*) is computed correctly on line 6.

Since the root of the gene tree can either be mapped to the unknown location or some actual syntenic region, we minimize over the two cases and get the optimal cost for reconciling the entire gene tree. Using standard DP traceback techniques, we can also obtain the mappings and events involved in an optimal solution.

